# Loss of PI3Kδ activity drives autoimmune colitis by impairing extrathymic Treg differentiation

**DOI:** 10.1101/2024.11.21.624633

**Authors:** Ee Lyn Lim, Yamin Qian, Fuminori Sugihara, Atsushi Tanaka, Shimon Sakaguchi

## Abstract

Peripherally-derived regulatory T cells (pTregs) have a prominent role in maintaining intestinal immune homeostasis. In cases of phosphoinositide-3-kinase δ (PI3Kδ) inactivation, such as in patients receiving PI3Kδ inhibitor idelalisib as a cancer treatment, breakdown of intestinal immune tolerance occurs frequently in the form of diarrhoea and colon inflammation. In a mouse model of systemic PI3Kδ inactivation, both enhancement of anti-tumor immunity and colitis have been described, as a result of Treg impairment. However, in view of the critical role for Tregs in the prevention of systemic autoimmunity, the basis for such tissue-restricted breach of immune tolerance upon loss of PI3Kδ function is not yet understood. We report here that mice lacking PI3Kδ activity do not suffer a general defect in Treg immunosuppression, but specifically fail to develop Helios^-^ pTregs in the colon. We demonstrate reduced extrathymic Treg induction, in vitro and in vivo, from naïve CD4^+^ T cells with inactive PI3Kδ, along with dysregulation of a tissue-resident phenotype. These results suggest a non-redundant role for PI3Kδ-dependent pTreg differentiation in maintaining tolerance to commensal microbial antigens in the gut.

## Introduction

The intestinal environment presents a particular tightrope challenge to immune regulation: the distinction between friend and foe among myriad microbial and dietary antigens. Multiple levels of safeguards deter aberrant inflammatory responses, from the physical separation of microbes from immune surveillance by the mucosal barrier (1), to the anti-inflammatory specialization of gut-resident myeloid subsets (2,3), and the production of tolerizing metabolites by both host cells and commensal bacteria along the intestinal tract (4–6). Bowel inflammation commonly results from perturbation of these safeguards by genetic, dietary, or therapeutic causes, whether in the form of microbiome disruption, epithelial ‘leakiness’, or incapacitation of immune regulatory mechanisms (7).

A central component in the maintenance of intestinal immune homeostasis is regulatory T cells (Tregs) – more specifically, peripherally-derived Tregs (pTregs) in which the immuno-suppressive Foxp3-driven transcriptional program is induced in mature CD4^+^ T cells by environmental cues beyond the thymus. Both the role of Tregs in preventing systemic autoimmunity (8,9), and the particular significance of pTregs within the gut environment (10), have been extensively established. While genetic ablation of pTregs, by deletion of the Foxp3 CNS1 enhancer region (10), has provided some insight into the distinct anatomical and functional niches occupied by thymic Tregs (tTregs) and pTregs, it remains challenging to interrogate these Treg populations – and indeed, their more specialized subsets (11,12) – in an unperturbed immune system, due to a lack of reliable distinguishing phenotypic markers. Helios, an Ikaros family zinc-finger transcription factor, has been proposed to mark Tregs of thymic origin (13), and remains the current standard of distinction in spite of reports of upregulation in activated pTregs (14). A deeper understanding of the various mechanisms underpinning Treg-mediated immune tolerance in different physiological contexts would bring significant benefit to human health, especially in the management of therapy-induced immune-related adverse events (irAEs) (15,16).

In cancer therapy, the phosphatidyl-inositide-3-kinase δ (PI3Kδ) inhibitors idelalisib (Zydelig, Gilead) and to duvelisib (Copiktra, Secura Bio) are in clinical use to treat chronic lymphocytic leukemia, small lymphocytic leukemia and follicular lymphoma, but with significant risk of severe, even fatal adverse effects, including diarrhoea, colitis and intestinal perforation (16–18). The PI3K pathway exerts broad effects on cell proliferation, differentiation and mobility; the δ isoform of kinase subunit p110 is restricted in expression to leukocytes, and its pharmacological inhibition thus confers immune-modulatory effects (19). PI3Kδ inactivation has shown considerable preclinical promise also in non-hematological tumors (20–22), and its broader application, for example in strategic combination with other immunotherapies (23), would be of immediate value. Anti-tumor immune enhancement with PI3Kδ inactivation is shown to stem from a preferential impairment of Tregs (24), directly connecting the intended efficacy and unintended adverse effects of the therapy. In both cases, the tissue-restricted nature of immune activation – whether in the tumor or in the colon – resulting from disruption of a systemically-important mode of immune regulation invites questions on the specific changes to Treg biology in the absence of PI3Kδ.

In this work, we utilised a spontaneously-occurring autoimmune colitis model in mice bearing a systemic kinase-inactivating D910A mutation in PI3Kδ (PI3Kδ^D910A^), to interrogate the specific alterations in Treg biology caused by loss of PI3Kδ function. We found PI3Kδ^D910A^ Treg suppression to be largely intact on a per-cell basis, but skewed sharply at the population level with the absence of Helios^-^ pTregs within the colon. In vitro and in vivo extrathymic Treg induction were both impaired in CD4^+^ T cells lacking PI3Kδ activity. We propose a novel and indispensable role for PI3Kδ in the development and persistence of a pTreg population capable of mediating intestinal immune homeostasis. This discovery lays the foundation for better phenotypic analysis and therapeutic strategy in the clinical use of PI3Kδ inhibition.

## Results

### PI3Kδ^D910A^ mice developed colon-restricted inflammation

We observed diarrhoea and colon inflammation in PI3Kδ^D910A^ mice bred in our facility. Thickening of colon tissue was evident (Fig. 1a) from onset, commonly around 17-22 weeks of age (Fig. 1b). Histological and immunofluorescence analyses (Fig. S1a, 1c) further revealed a structural breakdown of villi. Prior to diarrhoea onset, mice from 8-9 weeks of age with no symptoms of overt inflammation were found to have a single enlarged lymph node in the mesentery, corresponding to the colon/cecum-draining C1 lymph node (5) (Fig. 1d, S1b).

**Fig. 1.**
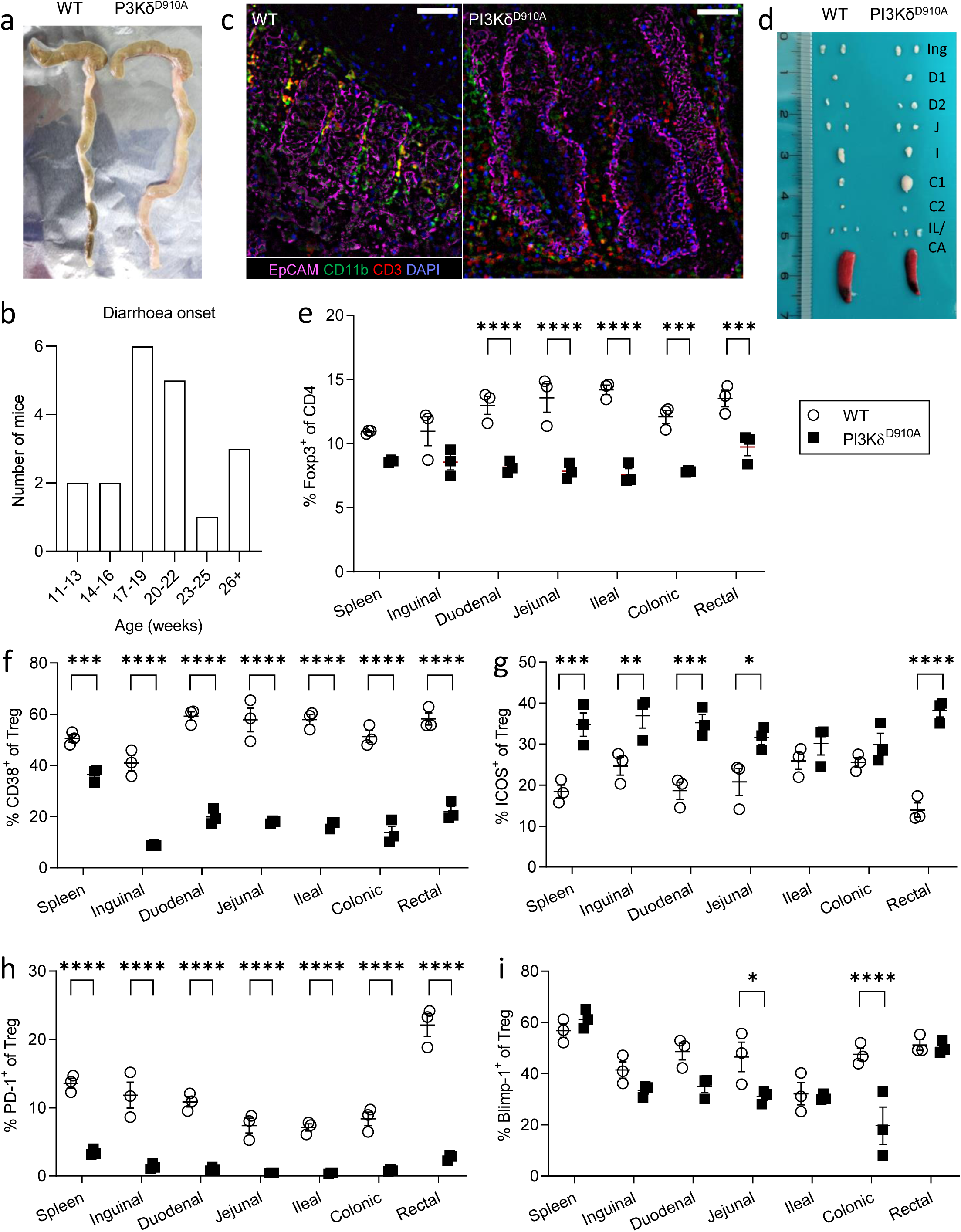
PI3Kδ^D910A^ mice develop colon-restricted inflammation and altered gut microbiome. (a) Macroscopic comparison of tissue thickening in PI3Kδ^D910A^ colon compared to age-matched WT. (b) Number of PI3Kδ^D910A^ mice which begun to develop diarrhoea at the specified age (total n = 19). (c) Immunofluorescent images of colon slices, showing epithelial cells (EpCAM), myeloid cells (CD11b), T cells (CD3), and nuclei (DAPI). Samples from age-matched WT and PI3Kδ^D910A^ mice. Scale bars indicate 100 μm. (d) Size comparison of dissected individual mLN, along with inguinal LN and spleen. (e) Quantification of Foxp3^+^ cells as a proportion of CD4^+^ T cells in spleen, inguinal LN and individual mLN. (f-i) Proportion of CD4^+^ Foxp3^+^ Tregs expressing CD38 (f), ICOS (g), PD-1 (h), and Blimp-1 (i). n = 3 biological replicates, representative of 2 independent experiments. *, p < 0.05; **, p < 0.01; ***, p < 0.001; ****, p < 0.0001.

In terms of cellular composition, we found a reduction in Tregs as a proportion of total CD4^+^ T cells in PI3Kδ^D910A^ mice compared to WT, in the draining lymph nodes of small intestine and colon (Fig. 1e). In the thymus, we reproduced previous reports of increased Foxp3^+^ CD4-single positive (CD4SP) cells (Fig. S1c) (25), and further observed a specific increase in CD25^+^ Foxp3^-^ Treg “precursor 1” cells over CD25^-^ Foxp3^+^ Treg “precursor 2” cells (Fig. S1b). PI3Kδ^D910A^ Tregs also showed consistently lower expression of CD38 (Fig. 1f), reported as a marker of high suppressive capacity (26), and elevated ICOS expression as previously shown (27), except in the draining lymph nodes of ileum and colon (Fig. 1g). PD-1 expression was strongly decreased in PI3Kδ^D910A^ Tregs (Fig. 1h). While PD-1 is thought to restrain Treg function and proliferation (28), its reduced expression in this instance may rather reflect a less activated state in PI3Kδ-deficient Tregs; CD8^+^ T cells and CD4^+^ Foxp3^-^ conventional T cells (Tconv), although expressing far lower levels of PD-1 at the resting state compared to Tregs, also showed reduced expression in PI3Kδ^D910A^ mice (Fig. S1d). Notably, Blimp-1, a transcription factor linked to the stability of the Treg immunosuppressive phenotype (29), was found to be specifically reduced in the mesenteric lymph nodes (mLN) of PI3Kδ^D910A^ mice compared to WT (Fig. 1i).

Both the development of inflammation in the colon and its remarkable specificity to the colon agree with the previously reported phenotype of PI3Kδ^D910A^ mice (30), albeit at greater severity than previously described. We note, however, that this mouse strain is completely asymptomatic in some facilities (31), suggesting that the breach in intestinal immune tolerance is driven by elements of the gut microbiome. 16s rRNA sequencing of fecal samples from pre-diarrhoeal mice (approx. 10 weeks old) revealed distinct clustering of PI3Kδ^D910A^ samples from WT controls by principal component analysis (PCA) (Fig. S2a, b). On the level of individual bacterial strains, we unexpectedly found no significant difference in the abundance of some known inflammation-promoting (e.g. *Prevotella spp.* (32,33), Fig. S2c) or dampening (e.g. family *Clostridiales* (34), Fig. S2d) strains, and other prominent strains were not detected at all (e.g. *Helicobacter spp.*). Only the *Bacteriodales* family *S24-7*, also known as *Muribaculaceae*, showed statistically significant reduction in PI3Kδ^D910A^ samples compared to WT (Fig. S2e). *Muribaculaceae* is a poorly-characterized component of both human and murine gut microbiota (35,36), but its proposed role as a protective commensal makes it an intriguing candidate for further investigation.

### PI3Kδ^D910A^ Tregs did not show a general loss of suppressive function

With previous reports of defects in Treg suppressive function upon PI3Kδ inactivation in the tumor immune response (20,21), we determined whether the colon inflammation in PI3Kδ^D910A^ mice was a consequence of fundamentally-impaired Treg suppression. Despite an overall reduction of Tregs compared to Tconv, we note that these mice did not develop systemic autoimmune inflammation, suggesting that overall immune homeostasis was maintained. To assay cell-intrinsic suppressive capacity, we measured expression of a key receptor in Treg suppression, CTLA-4, which blocks and removes costimulatory signals from APCs to effector T cells (37). Contrary to expectations, we found increased expression of CTLA-4 in PI3Kδ^D910A^ Tregs (Fig. 2a-c). A specific function of CTLA-4 in Tregs is the binding and removal of co-stimulatory ligands CD80 and CD86 from the surface of APCs by trogocytosis (38). Using an immortalized dendritic cell line JAWS, expressing GFP-fused CD80 and CD86, we quantified the trogocytic activity of PI3Kδ^D910A^ Tregs and found, in line with their elevated CTLA-4 expression, increased transfer of CD80- and CD86-GFP from the JAWS cells to the Tregs (Fig. 2d, e).

**Fig. 2.**
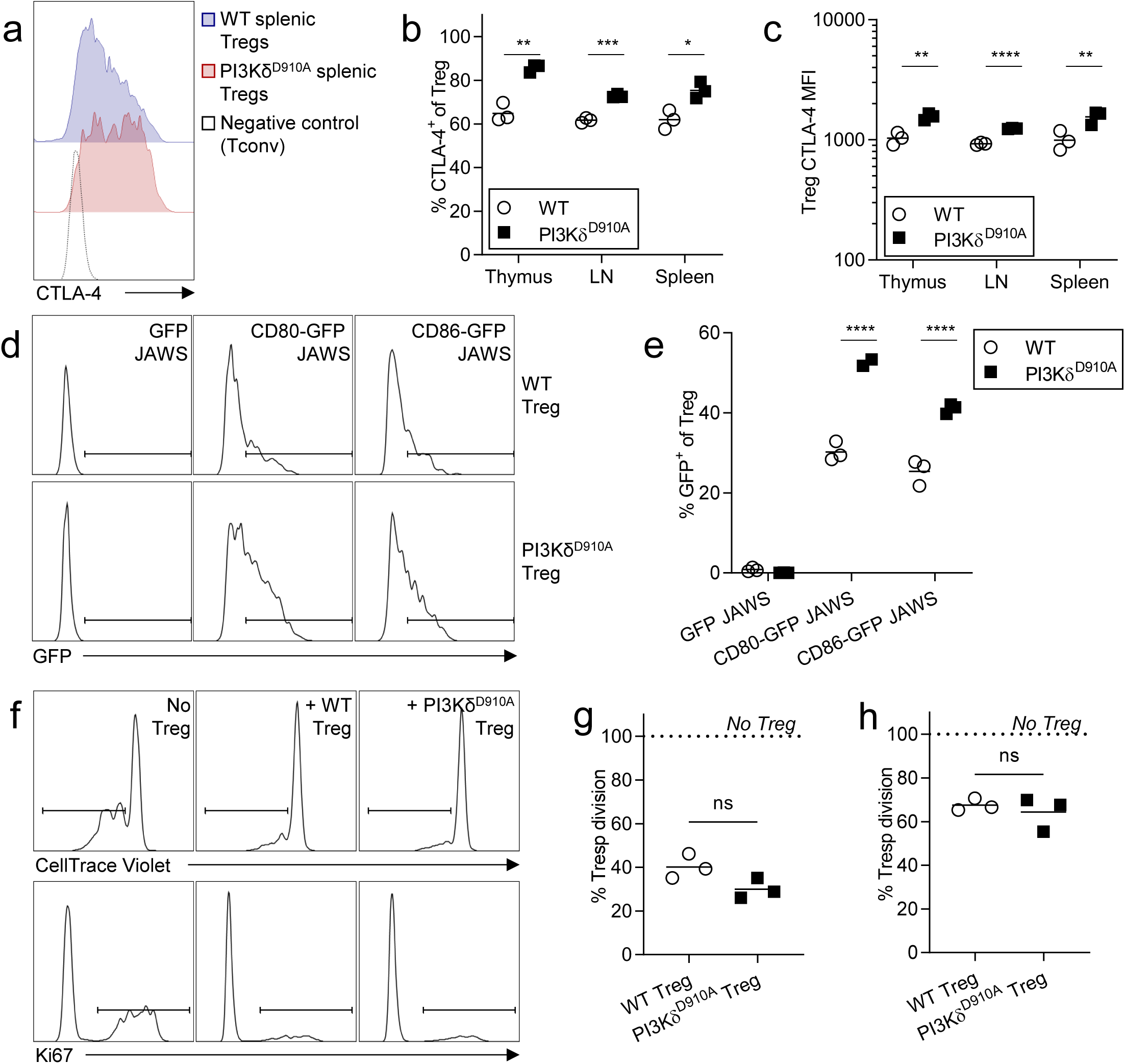
Tregs from PI3Kδ^D910A^ are not inherently defective in suppressive function. (a) Representative histograms of CTLA-4 expression, measured by flow cytometry. (b) Quantification of CTLA-4^+^ cells as a proportion of total Tregs. (c) Quantification of CTLA-4 median fluorescent intensity (MFI). n = 3 biological replicates. (d) Representative histograms of GFP trogocytosis by Tregs cocultured with JAWS DCs expressing GFP, CD80-GFP or CD86-GFP. (e) Quantification of GFP^+^ Tregs as a proportion of total Tregs in culture. n = 3 biological replicates, representative of 4 independent experiments. (f) Representative histograms of CellTrace Violet dilution or Ki67 expression in Tresp cultured with or without WT or PI3Kδ^D910A^ Tregs. (g-h) Quantification of proliferative Tresp stimulated with anti-CD3/CD28 Dynabeads (g) or anti-CD3-supplemented splenic DCs (h). n = 3 biological replicates. *, p < 0.05; **, p < 0.01, ***, p < 0.001; ****, p < 0.0001; ns, not significant.

To determine whether PI3Kδ^D910A^ Tregs suffer a cell-intrinsic impairment of suppressive capacity, we performed in vitro suppression assays to measure attenuation of conventional T cell proliferation. Using a cell-surface human CD2 (hCD2) reporter for Foxp3, CD4^+^ Foxp3-hCD2^+^ Tregs were sorted from WT and PI3Kδ^D910A^ mice and cocultured with WT CD4^+^ Foxp3-hCD2^-^ responder cells (Tresp) at a 1:2 ratio with TCR stimulation. Whether under stimulation with anti-CD3/CD28-coated Dynabeads (Fig. 2f, g) or with splenic dendritic cells (DCs) as antigen-presenting cells (APCs) (Fig. 2h), we found that PI3Kδ^D910A^ Tregs reduced Tresp proliferation to the same extent as WT Tregs.

These findings indicate that, on a per-cell basis, PI3Kδ^D910A^ Tregs are functionally normal or even slightly enhanced in suppressive function, suggesting a more context-specific driver for the colon inflammation.

### Immune phenotyping of colon infiltrate revealed specific absence of Helios^-^ Tregs

To identify an alternative explanation for the colon inflammation in the absence of general Treg impairment, we performed detailed phenotypic analysis, covering 30 lineage and functional markers (Table S3a) by mass cytometry, in WT and PI3Kδ^D910A^ mice pre- and post-inflammation onset (7 weeks and 22 weeks of age, respectively). In the intestinal tissue (including small intestine and colon intra-epithelial and lamina propria immune infiltrates) we found no reduction – and in some cases a slight increase – of Tregs in PI3Kδ^D910A^ mice compared to WT, in contrast to peripheral lymphoid tissues (Fig. S3b). UMAP dimensionality reduction of colon lamina propria (cLP) Tregs revealed a striking separation of WT and PI3Kδ^D910A^ cells from 22-week-old mice (Fig. 3a) – driven in part by a difference in the expression of Helios between the two cell populations (Fig. 3b, see Fig. S3c for complete expression data). A side-by-side comparison of Helios expression between cells from each mouse group confirms a near-complete absence of Helios^-^ Tregs in the cLP of 22-week-old PI3Kδ^D910A^ mice (Fig. 3c).

**Fig. 3.**
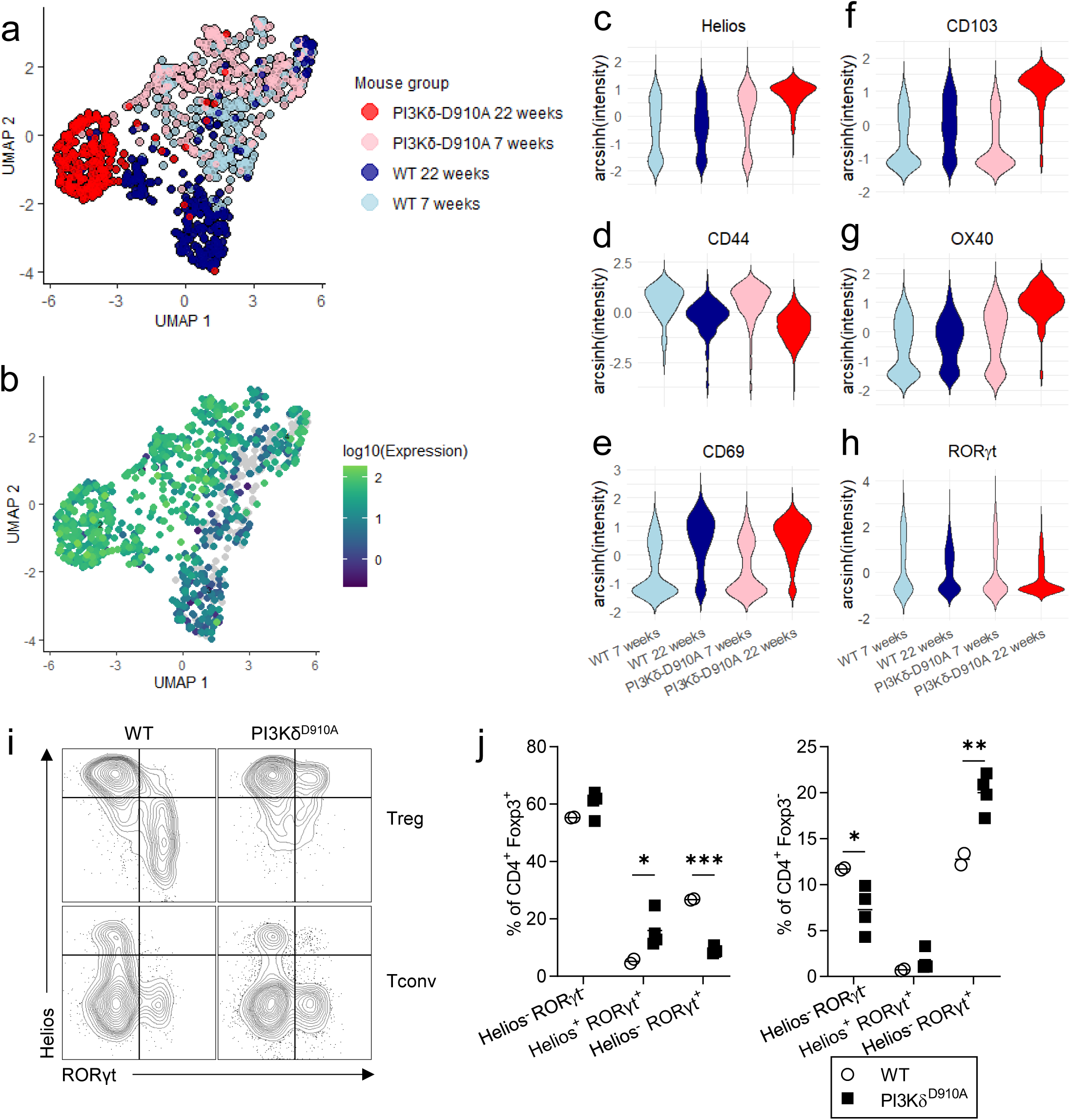
Colon Tregs from PI3Kδ^D910A^ mice highly express tTreg marker Helios. (a) UMAP dimensionality reduction on mass cytometric characterization of cLP Tregs from WT and PI3Kδ^D910A^ mice, before (7 weeks) and after inflammation onset (22 weeks). (b-c) Helios expression overlaid on UMAP (b) or in cells separated by mouse group (c). (d-h) Expression of activation and functional markers CD44 (d), CD69 (e), CD103 (f), OX40 (g), and RORγt (h) by mouse group. Mass cytometric data (a-h) pooled from n = 2 biological replicates for each group. (i-j) Flow cytometric measurement of Helios and RORγt expression in cLP Tregs and Tconv in post-inflammation PI3Kδ^D910A^ mice compared to age-matched WT (i, representative plots; j, quantification). n = 2 (WT) or 4 (PI3Kδ^D910A^) biological replicates. *, p < 0.05; **, p < 0.01, ***, p < 0.001.

To assess the possibility that Helios expression can also indicate recently-activated pTregs (14) – a definite consideration in the inflamed colon of older PI3Kδ^D910A^ mice – we showed that T cell activation markers CD44 and CD69, with the latter especially indicating recent TCR engagement, were not increased in PI3Kδ^D910A^ cLP Tregs compared to WT cells from mice of the same age (Fig. 3d, e). Other markers strongly elevated in post-inflammation onset PI3Kδ^D910A^ cLP Tregs were CD103, an integrin widely reported to be important for Treg suppression in tissues including the colon (39) (Fig. 3f), and OX40, a TNF receptor super-family member which can restrict Treg suppressive function upon ligand engagement (40) (Fig. 3g); whether the high expression of these markers are a cause or effect of the breach in colon immune homeostasis in PI3Kδ^D910A^ mice remains to be determined.

pTregs with a Helios^-^ RORγt^+^ phenotype have been particularly implicated in maintaining intestinal immune tolerance (41). Despite the nearly uniform expression of Helios in PI3Kδ^D910A^ cLP Tregs as observed in mass cytometry data, RORγt expression on these same cells was not markedly different between WT and PI3Kδ^D910A^ mice (Fig. 3h). We confirmed this result by flow cytometric analysis on a separate cohort of WT and PI3Kδ^D910A^ mice aged 22-23 weeks, demonstrating the loss of the Helios^-^ RORγt^+^ Treg subset and a concomitant increase in a Helios^+^ RORγt^+^ population in PI3Kδ^D910A^ cLP (Fig. 3i, j). We also observed an increase in RORγt^+^ Tconv in the colon of PI3Kδ^D910A^ mice, consistent with an accumulation of Th17 cells (Fig. 3i, j) – an effector T cell population frequently implicated in inflammation, and most significantly in irAEs upon PI3Kδ inhibition (16).

These findings demonstrate a specific deficiency of Helios^-^ pTregs in PI3Kδ^D910A^ mice, potentially compensated for in numbers by increased accumulation of tTregs, which nonetheless fail to replicate pTreg functions in intestinal immune homeostasis.

### In vitro-differentiated iTregs show impaired upregulation of key suppressive markers

Considering difficulties in reliably distinguishing tTregs and pTregs in vivo, we turned to an in vitro model of TGF-β-driven extrathymic Treg induction. Naïve CD4^+^ Foxp3-hCD2^-^ T cells were cultured with TCR stimulation from plate-coated anti-CD3/anti-CD28, combined with IL-2, TGF-β and antibody blockade of cytokines driving T helper differentiation (IL-4, IL-6, IL-12 and IFN-γ). Differentiation of Foxp3^+^ induced Tregs (iTreg) was drastically reduced in PI3Kδ^D910A^ cultures throughout the 6-day experimental duration (Fig. 4a). This difference persisted across a range of TGF-β concentrations, as measured after 4 days in culture (Fig. 4b). Intriguingly, replacing plate-bound antibody stimulation with APC stimulation (B cells supplemented with anti-CD3) significantly closed the gap between WT and PI3Kδ^D910A^ iTreg induction, yielding a near-equal proportion of Foxp3^+^hCD2^+^ cell from both cultures despite an initial delay (days 1-3) in Foxp3 expression in PI3Kδ^D910A^ cells (Fig. 4c). The difference between these methods of stimulation point to the contributions of complex costimulatory interactions, beyond CD28 engagement, provided by APCs, which can partially compensate for a weakened TCR signal due to PI3Kδ inactivation.

**Fig. 4.**
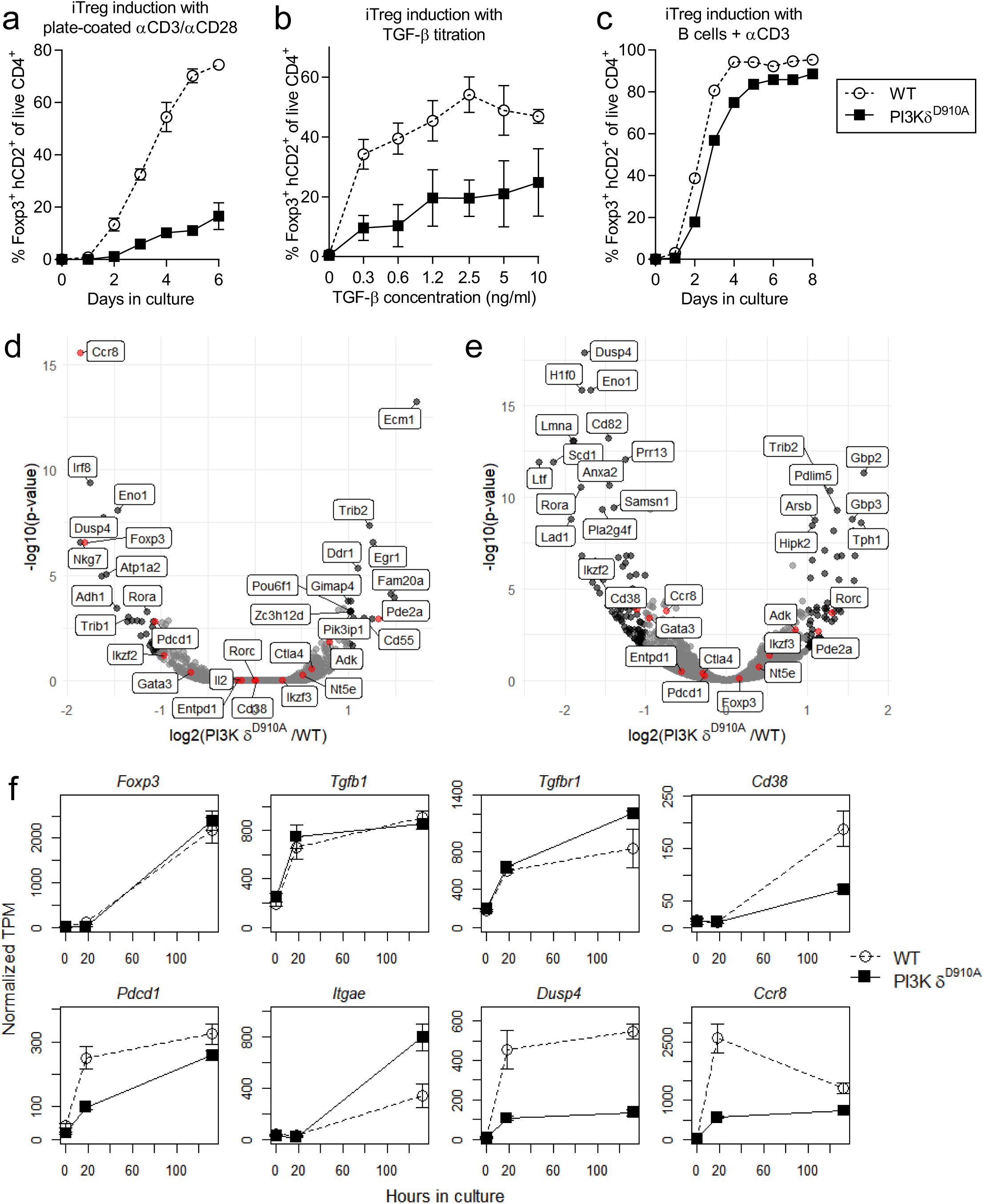
In vitro-differentiated iTregs reveal transcriptomic and proteomic differences. (a) Time course of Foxp3 induction in naïve CD4^+^ Foxp3^-^ T cells cultured with TGF-β and stimulation with plate-coated anti-CD3 and anti-CD28. n = 3, representative of 2 independent experiments. (b) Titration of TGF-β concentration versus Foxp3 induction after 4 days in culture. n = 4 combined from 4 independent experiments. (c) Time course of Foxp3 induction in naïve CD4^+^ Foxp3^-^ T cells cultured with TGF-β and stimulation with anti-CD3-supplemented B cells. n = 3, representative of 4 independent experiments. (d-e) RNA sequencing of WT and PI3Kδ^D910A^ iTregs induced with TGF-β, after 18 hours (sorted for Foxp3-hCD2^-^, a) or 5 days (sorted for Foxp3-hCD2^+^, b) in culture. (f) Time course of expression changes in selected genes over the duration of iTreg culture. Genes marked in black show >2-fold change with adjusted p-value < 0.05 in PI3Kδ^D910A^ cells compared to WT; genes marked in red indicate known roles to Treg development or function. n = 3 biological replicates.

Using B cell-stimulated iTregs to obtain requisite cell numbers from both WT and PI3Kδ^D910A^ samples, we performed transcriptomic analysis at two time points in iTreg differentiation – firstly, on Foxp3-hCD2^-^ cells after 18 hours in culture, to quantify transcriptional changes directly following TCR stimulation and preceding Foxp3 expression (Fig. 4d); secondly, on Foxp3-hCD2^+^ cells after 5 days in culture, to interrogate mature iTregs (Fig. 4e). Together with transcriptional profiling of ex vivo naïve CD4^+^ Foxp3-hCD2^-^ T cells, we constructed a broad timeline of differential gene expression in WT vs PI3Kδ^D910A^ iTregs (Fig. 4f).

Even in cells sorted for the absence of Foxp3 reporter hCD2, WT cells had slightly increased Foxp3 mRNA compared to PI3Kδ^D910A^, reflecting greater induction efficiency early in culture (Fig. 4c, d, f). Neither TGF-β nor its receptor (*Tgfb1* and *Tgfbr1*) were reduced at the transcriptional level in PI3Kδ^D910A^ iTregs compared to WT (Fig. 4f), suggesting a defect in downstream signal transduction. Consistent with observations in ex vivo cells, CD38 and PD-1 expression remained low in PI3Kδ^D910A^ iTregs throughout the course of stimulation (Fig. 4f, confirmed at protein level in Fig. S4a).

As in colonic Tregs, we found increased CD103 (*Itgae*) expression in PI3Kδ-inactivated iTregs, a marker for tissue infiltration present even in the in vitro setting (Fig. 3f, 4f). On the other hand, the dual-specificity phosphatase 4 (Dusp4), reported to specifically promote expression of activation- or tissue residence-related genes in Tregs (42), is rapidly upregulated in WT iTregs upon stimulation, but no response is observed in PI3Kδ^D910A^ iTregs (Fig. 4f). Similarly, the chemokine receptor CCR8 is sharply upregulated in WT cells after 18 hours in culture, but this transcriptional change does not occur in PI3Kδ^D910A^ cells, where expression remains significantly lower than WT even after a decline in expression in WT cells after 5 days (Fig. 4d-f). Proteomic analysis of Foxp3-hCD2^+^ iTregs after 4 days of culture confirmed differential expression of CCR8 (Fig. S4a). CCR8 has significance as a marker of highly-suppressive tissue-infiltrating Tregs, especially in the context of tumor immunosuppression (43,44). We hypothesize that reduced CCR8 expression in PI3Kδ^D910A^ extrathymically-differentiated Tregs could impair their ability to infiltrate or reside in the colon, but this proved difficult to demonstrate in vivo: cLP Tregs in both WT and PI3Kδ^D910A^ mice were almost 100% CCR8^+^ (Fig. S4b, c), suggesting that CCR8 expression is required for gut tissue infiltration. Paradoxically, we found a larger proportion of CCR8^+^ Tregs in the spleens of PI3Kδ^D910A^ compared to WT, consistent with a compensatory, but functionally insufficient increase in tTreg activation.

In summary, the defects in iTreg induction and the differences in gene expression between WT and PI3Kδ^D910A^ iTregs reflect perturbations in TCR signaling, reduced responsiveness to TGF-β, and disruption of tissue infiltration and persistence, that offer insight into the loss of pTreg-mediated tolerance in the colon.

### In vivo extrathymic Foxp3 induction is impaired in the absence of PI3Kδ activity

To directly assay extrathymic Treg differentiation in vivo, we measured pTreg induction from congenically marked, naïve Foxp3^-^ CD4^+^ T cells, adoptively transferred into RAG2^-/-^ hosts. To minimize complications from potential T cell transfer-induced inflammation, WT Foxp3^+^ CD4^+^ Tregs were co-transferred in a 1:2 ratio with the naïve Tconv to ensure a non-colitogenic experimental setting. 5 weeks after transfer, spleen, mLN, small intestine, and colon tissue were recovered from host mice and analysed by flow cytometry. Foxp3 expression among PI3Kδ^D910A^ donor cells was much reduced compared to WT in all tissues, and especially in the cLP (Fig. 5a, b), mirroring impairments in in vitro differentiation.

**Fig. 5.**
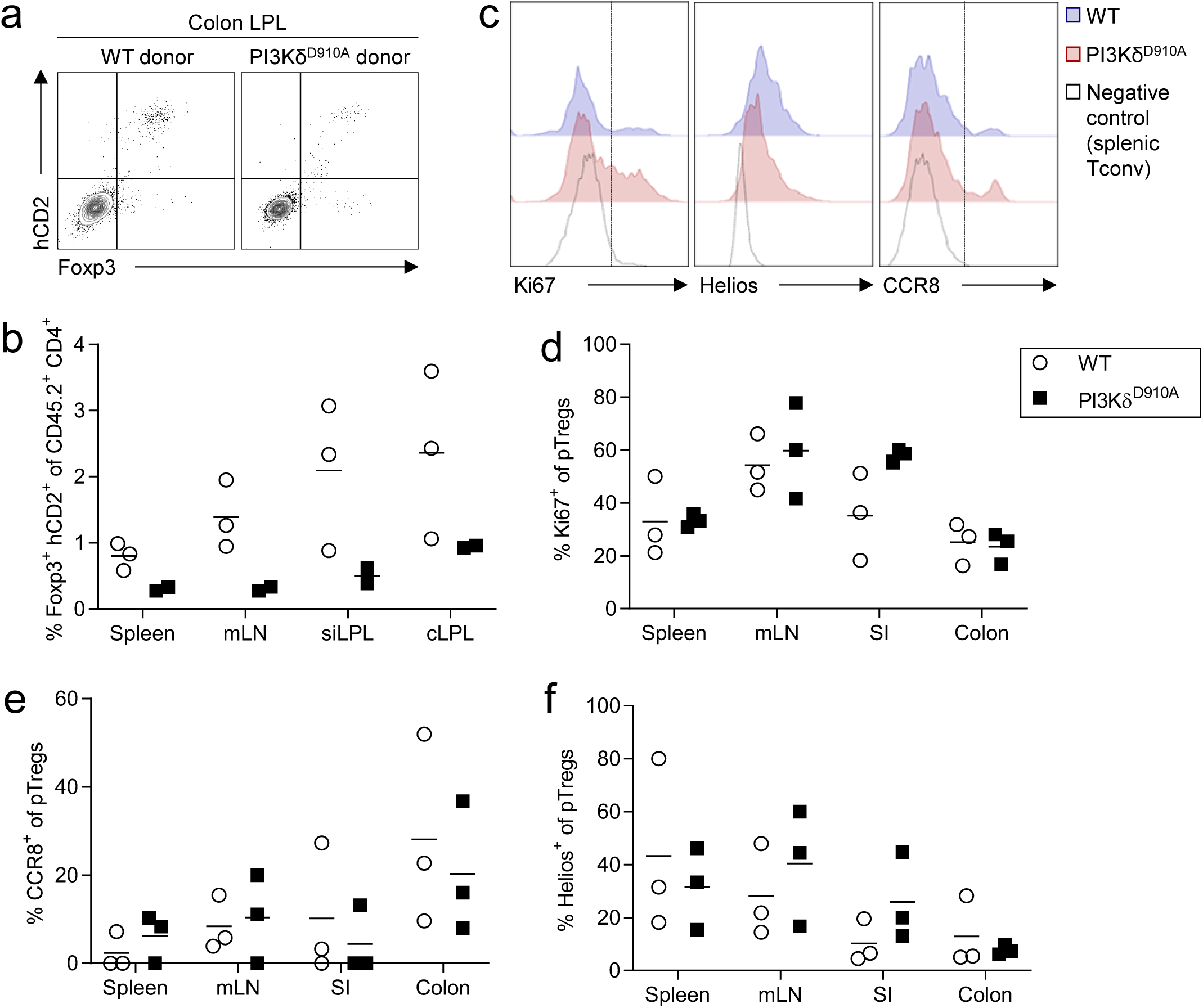
Extrathymic Treg differentiation is impaired in the absence of PI3Kδ activity. (a-b) Foxp3 expression among CD45.2^+^ CD4^+^ Foxp3-hCD2^-^ T cells transferred into CD45.1^+^ RAG2^-/-^ hosts 5 weeks post-transfer (a, representative plots from cLP; b, quantification). n = 2 or 3 biological replicates, representative of 3 independent experiments. (c-f) Ki67, Helios and CCR8 expression in pTregs measured by flow cytometry (f, representative plots; g-i, quantification). n = 3 biological replicates.

The adoptive transfer assay provided an opportunity to phenotypically interrogate a pure population of experimentally-derived pTregs. We found equal or slightly elevated expression of Ki67 in PI3Kδ^D910A^ pTregs, excluding a proliferation defect as the cause of reduced Foxp3^+^ cell numbers (Fig. 5c, d). CCR8 expression was also not different between WT and PI3Kδ^D910A^ cells (Fig. 5c, e), but was overall far lower than that observed in endogenous colon-infiltrating Tregs (Fig. S4b, c), perhaps as a consequence of the RAG2^-/-^ immunodeficient context. Helios expression was low in both WT and PI3Kδ^D910A^ pTregs, especially in the colon (Fig. 5c, f), in contrast with endogenous Tregs in the PI3Kδ^D910A^ colon (Fig. 3c), demonstrating that Helios upregulation is not an intrinsic trait of PI3Kδ inactivation in Tregs.

Taken together, these results provide direct evidence that PI3Kδ-inactivated CD4^+^ T cells have reduced capacity to differentiate into pTregs in vivo.

## Discussion

This work establishes a critical role for PI3Kδ activity in TGF-β-mediated extrathymic Treg induction, in mediating the combined TCR and TGF-β signals required for Foxp3 expression. In a mouse model of systemic PI3Kδ inactivation, we report the loss of a colon-resident population of Helios^-^ Tregs, and impaired differentiation of Foxp3^+^ iTregs in vitro and pTregs in vivo. We also show a disrupted tissue-resident phenotype, exemplified by reduced expression of the chemokine receptor CCR8 in PI3Kδ-deficient extrathymic Tregs, suggesting a further incapacity to infiltrate gut tissues.

We note that the original description of PI3Kδ-inactivated Tregs found a defect in the ability to suppress Tresp stimulated with APCs or CD3/CD28 Dynabeads (25), which differs from findings presented here (Fig. 2b, c). Subtle differences in the experimental setup which may explain the discrepancy are the use of T cell-depleted splenocytes as APCs and a ratio of 1 Dynabead:5 Tresp in the original report, compared to purified splenic DCs and a ratio of 1 Dynabead:2 Tresp in this study – in both cases we have applied a more potent stimulus, which may be sufficient to fulfil a higher requirement to elicit suppression from PI3Kδ-deficient Tregs.

Dysregulation of the T cell tissue infiltration capacity delineates two distinct parts in the impact of PI3Kδ inactivation on Treg-mediated intestinal immune homeostasis. In vivo, colon-infiltrating Tregs in mice universally express CCR8, indicating a requirement for CCR8 for their localization to and persistence in the colon. Extrathymic Treg differentiation is impaired by a lack of PI3Kδ function, and the pTregs which do develop in PI3Kδ-deficient mice further fail to upregulate CCR8 and Dusp4, and do not populate the intestinal niche. On the other hand, CCR8-expressing Helios^+^ tTreg numbers increase in the colon, perhaps in compensation, but are unable to control microbiome-triggered inflammation. We observe even a paradoxical increase in CCR8^+^ Tregs in the spleens of PI3Kδ^D910A^ mice (Fig. S4b, c), raising the possibility that tTregs populating tissue niches left vacant by the absence of pTregs fail to persist in those environments, and are purged back into circulation.

The confined effects of PI3Kδ inactivation to pTreg induction, leaving intact the major suppressive functions of the bulk of the Treg population, draws intriguing connections with previous studies on PI3Kδ inhibition as a cancer immunotherapy (20,21). Definitive proof for the role of pTregs in tumor immunosuppression has proven elusive, not least because of a lack of reliable markers to distinguish endogenous tTregs and pTregs, and the vast variability of the immune response between tumor types. Considering the relatively late age of onset of the colon inflammation in PI3Kδ^D910A^ mice, and its presence or absence in the context of facility-dependent microbial milieus, we hypothesize that the pTreg population lost from PI3Kδ-deficient mice may respond specifically to one or more commensal microbial antigens – recognized as foreign, but tolerated, in an unperturbed T cell repertoire. This forms a strong parallel with tumor neo-antigens, which are not presented as part of ‘self’ in the thymic development of central tolerance, but are enforced as such by a tolerizing tumor microenvironment.

A third context in which tolerance to foreign antigen is crucial to immune homeostasis emerges cancer patients treated with idelalisib, among whom transaminitis from liver tissue damage, alongside colon inflammation, is a prominent adverse effect (16,45). More than in specific pathogen-free laboratory mice, human livers are tasked with clearing an array of environmental detritus from the blood without triggering an inflammatory response, and are hence susceptible to immune dysregulation when pTreg function is compromised. Understanding the source of these adverse effects in patients would be pivotal to the further development of PI3Kδ inhibitors in clinical therapy.

A key question that remains outstanding is the mechanistic link between a loss of PI3Kδ activity and the demonstrated defects in extrathymic Treg differentiation. While it is known that TCR signaling is attenuated in the absence of functional PI3Kδ (30), we show here that costimulatory cues provided by APCs can largely rescue the Foxp3 induction impairment in vitro (Fig. 5a vs b). Which specific costimulatory interaction plays this role, whether it is present in gut environment, and how PI3Kδ is involved in transducing its signal, are subjects for future investigation.

Moreover, a weakened TCR signal in itself can have far-reaching effects on the T cell repertoire as a whole, as we recently reported (46), shifting the range of TCR specificities available to respond to self- and non-self-antigens. With PI3Kδ inactivation, TCRs recognizing commensal antigens may no longer signal strongly enough to trigger the Foxp3 transcriptional program, producing instead a population of microbiome-reactive conventional T cells. Even among developing thymic Tregs, our findings suggest a relative increase in the contribution of CD25^+^ Foxp3^-^ precursor 1 cells over CD25^-^ Foxp3^+^ precursor 2 cells to the mature Treg pool in PI3Kδ-inactivated mice (Fig. S1c). These precursor populations are reported to have distinct TCR repertoires – with a lower affinity among precursor 2 cells for thymic self-antigens – and distinct abilities to mediate immunosuppression in different inflammatory contexts (47). The functional divergence between Treg subsets, whether thymic/peripheral (48) or precursor 1/precursor 2-derived (47), lends credence to our postulation that compensatory infiltration and activation among PI3Kδ^D910A^ tTregs cannot fill the same niche as the missing pTregs.

We also show reduced responsiveness of PI3Kδ-inactivated naïve CD4^+^ T cells to TGF-β in culture (Fig. 4b), without a reduction in the expression of the TGF-β receptor (Fig. 4f, S4a). Direct engagement of the PI3K pathway downstream of TGF-β receptor binding is not widely established, especially in the context of immune cells, but in tumor cells, where both PI3K and TGF-β are strong drivers of transformation and proliferation, multiple points of cross-talk between the two pathways have been reported (49–51). In one specific example, the serine/threonine kinase PDK1, activated by PI3K, can phosphorylate and inactivate SMAD7, the inhibitory component of the TGF-β pathway (52,53), implying that PI3K inactivation may in turn attenuate the TGF-β signal. The exact nature of the interactions between these pathways likely depends on the broader cellular context, and the molecular chain of events within PI3Kδ-deficient naïve CD4^+^ T cells receiving a TGF-β signal remains to be elucidated.

## Acknowledgements

We thank Prof. Klaus Okkenhaug for the provision of PI3Kδ^D910A^ mice and critical reading of the manuscript, Prof. Shohei Hori for the provision of Foxp3-hCD2 mice, Dr. Wai Tuck Soh and Dr. Andrea Graziadei for advice on data analysis, Dr. Atsushi Sugimoto and Mr. Masaya Arai for technical instruction, and Ms. Kazumi Ishihara and Ms. Yamami Nakamura for technical assistance. ELL was supported by an IFReC Advanced Postdoctoral Fellowship research grant. This work was supported in part by JSPS KAKENHI Grant-in-aid 19K16692 (to ELL), 17K15723, 22H02920 (to AT), grants-in-aid from the Ministry of Education, Sports, and Culture of Japan 16H06295 (to SS), Japan Agency for Medical Research and Development P-CREATE 18cm0106303, LEAP 18gm0010005 (to SS), and P-PROMOTE 23ama221319 (to AT and SS). The authors declare no conflicting financial interests.

## Methods

### Mice and tissues

All mice used in this study were on the C57BL/6J genetic background, maintained under specific pathogen-free conditions in the experimental animal facilities at the Immunology Frontier Research Center, Osaka University, in accordance with institute regulations. Animal experiment protocols were approved by the institutional review boards of Osaka University. C57BL/6J mice were purchased from CLEA Japan. p110δ^D910A/D910A^ (PI3Kδ^D910A^) mice (30), with a homozygous kinase-inactivating aspartic acid–to-alanine (D910A) point mutation in the catalytic domain of p110δ, were provided courtesy of Prof. Klaus Okkenhaug, University of Cambridge, with permission from the Ludwig Institute for Cancer Research. Foxp3^tm1(CD2/CD52)Shori^ (Foxp3-hCD2) mice (54), which express human CD2 as a cell surface reporter for Foxp3, were provided courtesy of Prof. Shohei Hori, University of Tokyo. CD45.1 RAG2^-/-^ mice were previously described (55).

### Tissue homogenization

Single cell suspensions were obtained from spleens and LN by mechanical homogenization through a 30-70 μm filter mesh in PBS. Splenic cells were further treated with red blood cell lysis buffer (Sigma) for 5 mins at RT, then washed with PBS with 2% FBS.

Small intestines (dissected between stomach and cecum, Peyer’s patches removed from WT and absent from PI3Kδ^D910A^ mice) and colons (from cecum to rectum) were cut lengthwise and washed 3-4x in PBS to remove mucus and fecal matter. Cleaned tissues were cut into ∼2cm pieces, placed into RPMI 1640 with 0.5 mM EDTA and 2% FBS and agitated vigorously for 1 minute. Supernatant was collected (intraepithelial (IE) fraction) and combined with that from a second agitation. Tissue pieces were further finely minced (∼0.5 cm) and incubated in RPMI 1640 with 0.5 mg/ml collagenase D (Roche) and 2% FBS, with continuous stirring at 37°C for 2x 20 minutes. Supernatant was filtered through a 70 μm filter mesh, and remaining solid tissue pressed through with a syringe plunger (LP fraction).

Both IE and LP fractions were pelleted, resuspended in 40% Percoll (Cytiva) and layered over 80% Percoll, then centrifuged at 600 xg for 20 minutes at 20°C with no acceleration or brake. Leukocytes at the density interface were collected and washed with PBS with 2% FBS.

### Flow cytometry and cell sorting

A full list of antibodies and flow cytometric reagents used in this study can be found in Supplemental Information (SI). In brief, cell surface staining was performed in Brilliant Staining Buffer (BD Horizon) for 30 minutes on ice. In analysis or sorting of live cells, dead cell exclusion was achieved by the addition of 4’,6-diamidino-2-phenylindole (DAPI, Sigma-Aldrich) immediately before acquisition. For analysis of fixed cells, LIVE/DEAD™ Fixable Near-IR Dead Cell Stain (Thermo Fisher Scientific) was added with surface-staining antibodies, followed by fixation and permeabilization with Foxp3/Transcription Factor Staining Buffer Set (eBioscience) prior to intracellular staining, in permeabilization buffer for 30-60 minutes on ice. Wash steps were performed with PBS supplemented with 2% FCS. UltraComp eBeads (Invitrogen) were used as compensation controls. Cell sorting was performed on BD FACSAria sorters. Data acquisition was performed on BD LSRFortessa and FACSCanto analysers. Data analysis was performed in FlowJo (BD).

### Mass cytometry

Complete information regarding antibodies and reagents used in mass cytometry experiments is included in SI. Samples were acquired on a Helios mass cytometer (Standard Biotools). Data debarcoding and cell subset gating was performed in Cytobank Premium (Beckman Coulter). Data clustering (k-means), partitioning (Leiden) and dimensionality reduction (UMAP) was performed with the Monocle3 package in R (56–59).

### Immunohistochemistry and immunofluorescence imaging

Colon and small intestine sections were fixed in 10% paraformaldehyde and further processed by the Histopathology Core Facility, Niigata University Faculty of Medicine. Tissue sections were embedded in paraffin and slices stained with hematoxylin and eosin (HE). Images of the slides were then acquired on a Keyence BZ-X700 microscope with a bright-field filter.

For immunofluorescence staining, colon sections were embedded in low-melting point agarose and sliced at 300 μm thickness with a vibratome in a cold PBS bath. Slices were fixed for 2 hours with 4% PFA in PBS, then stained at 4°C overnight in anti-Ep-Cam APC (G8.8), anti-CD11b FITC (M1/70), anti-CD3 PE (145-2C11, all antibodies from Biolegend) and DAPI in PBS supplemented with 2% FCS. After washing, slices were mounted in glass-bottom dishes and imaged on a Keyence BZ-X700 microscope with 10x magnification.

### Suppression assay

Splenic DCs were first enriched from WT splenocytes using MACS CD11c microbeads, mouse, and MACS LS columns (Miltenyi Biotec) then sorted (CD11c^+^ DAPI^-^). Naïve CD4^+^ T cells (Tresp, CD4^+^ CD25^-^ CD44^-^ CD62L^hi^ DAPI^-^) were sorted from the spleen and LN of CD45.1 WT mice, then incubated in 1 μM CellTrace Violet (Thermo Fisher Scientific) in unsupplemented RPMI medium for 10 minutes at 37°C. Staining was quenched with cold RPMI medium containing 10% FBS and cells washed 3x with cold FBS-supplemented medium. Tregs (CD4^+^ hCD2^+^ CD25^+^ DAPI^-^) were sorted from the spleen and LN (after B cell depletion with MACS CD19 microbeads, Miltenyi Biotec) of CD45.2 Foxp3-hCD2 WT and PI3Kδ^D910A^ mice. Tresp were cocultured in 96-well U-bottom plates with or without Tregs at a ratio of 2:1, with the addition of either DCs (1 cell:3 Tresp) with 0.5 μg/ml purified anti-CD3 (145-2C11, Biolegend), or Dynabeads Mouse T-Activator CD3/CD28 for T-Cell Expansion and Activation (Gibco, 1 bead:2 Tresp), in RPMI 1640 medium (Nacalai Tesque) supplemented with 10% FBS. Cell proliferation was assessed by CellTrace Violet dilution or Ki67 staining after 4 days.

### Trogocytosis assay

Tregs (CD4^+^ CD25^+^ GITR^+^ DAPI^-^) were sorted from the spleen and LN (after B cell depletion) of WT and PI3Kδ^D910A^ mice, and cocultured overnight with JAWS immortalized dendritic cells expressing GFP or fusion constructs of CD80-GFP and CD86-GFP (38). Transfer of GFP to Tregs was assess by flow cytometry.

### iTreg induction assay

B cells (CD19^+^ DAPI^-^) were sorted from spleens of CD45.1 mice. Naïve CD4^+^ T cells (CD4^+^ hCD2^-^ CD25^-^ CD44^-^ CD62L^hi^ DAPI^-^) were sorted from spleens and LN (after B cell depletion) of CD45.2 Foxp3-hCD2 WT and PI3Kδ^D910A^ mice, and cocultured with B cells at a ratio of 2:1 in in RPMI 1640 medium supplemented with 10% FBS, 100 IU/ml recombinant human IL-2 (Shionogi), recombinant murine TGF-β (R&D) at 10 ng/ml unless otherwise stated, and anti-IL-4 (11B11, Invitrogen), anti-IL-6 (MP5-20F3, Biolegend), anti-IL-12/p40 (C17.8, eBioscience) and anti-IFNγ (XMG1.2, eBioscience), each at 1 μM. For Foxp3 induction time courses, cells were removed from culture every 24 hours and assessed for hCD2 and Foxp3 expression by flow cytometry. For TGF-β titration and proteomic characterization, cells were assessed after 4 days in culture. For transcriptomic analysis, cells were sorted for CD4^+^ hCD2^-^ DAPI^-^ after 18 hours and CD4^+^ hCD2^+^ DAPI^-^ after 5 days.

### RNA sequencing

16S rRNA metagenomic sequencing was carried out on frozen fecal pellets by the NGS facility at the Research Institute for Microbial Diseases, Osaka University. For transcriptomic sequencing, cells were lysed with RLT buffer (Qiagen) with 2-mercaptoethanol (Sigma-Aldrich) and RNA was purified using Agencourt RNAClean XP (Beckman Coulter). cDNA synthesis and amplification was performed using SMART-Seq v4 Ultra Low Input RNA Kit for Sequencing (Clontech), followed by fragmentation with Covaris Focused-ultrasonicator S220 (Covaris). The sequence library was constructed using KAPA Hyper Prep Kit (KAPA Biosystems) and sequenced on Ion S5 XL (Thermo Fisher Scientific) with single-end reads. Quality control, trimming, alignment and read quantification were respectively performed with the FASTQC, Trimmomatic, HISAT2 and Kallisto Quant tools in Galaxy (usegalaxy.org). Statistical analysis of differentially-expressed genes was determined via DESeq2 (Bioconductor R package (60,61)).

### Mass spectrometry protein identification

The protein solution from the cell samples were prepared by beads crasher with glass beads in PTS solution (62). The protein solutions were processed by trypsin after after treated with TCEP and iodoacetamide. The processed protein solution was cleaned up by C18 tip (GL-science) and quantified by fluorescamine and fluorescence analyzer (NanoDrop3300, Thermo). Then, the samples were fractionated by using SDB-SCX column tip (GL science) with TFA acidic condition (63).

The fractionated sample solutions were applied for data independent analysis (DIA) proteomics method of TIMS TOF Pro with nanoElute (Bruker). The mobile phase A: 0.1% FA water, B: 0,1% FA acetonitrile (Fujifilm Wako), the column was Nikkyo NTCC360 column (75um i.d., 12cm length). column temperature was kept 50oC, elution condition was 0%B to 35%B in 40min. The acquired data were analyzed by DIA-NN (64) with mouse protein database (Uniprot).

### Adoptive transfer pTreg induction

Naïve T cells were sorted as above from CD45.2 Foxp3-hCD2 WT and PI3Kδ^D910A^ mice. Tregs (CD4^+^ CD25^+^ GITR^+^ DAPI^-^) were sorted (after B cell depletion) from the spleens of CD45.1 WT mice. 10^6^ naïve T cells from each donor were mixed with 5x10^5^ WT Tregs and transferred by intravenous injection into 1 CD45.1 RAG2^-/-^ recipients. After 5 weeks, spleens, LN and intestinal tissue were collected from recipients and analysed by flow cytometry for the presence of donor-derived Tregs (CD45.2^+^ Foxp3^+^ hCD2^+^).

### Statistical analysis and data visualization

Statistical analysis was performed in GraphPad Prism using non-parametric (Mann-Whitney) t tests for data without multiple comparisons, and two-way ANOVA with Holm-Sidak correction for data with multiple comparisons. Data transformation and visualization was performed in R (65) using the packages tidyverse (66), dplyr (67), tibble (68), bestNormalize (69), gplots (70), ggrepel (71) and RColorBrewer (72).

