## Supplementary figures for "Loss of PI3Kδ activity drives autoimmune colitis by impairing extrathymic Treg differentiation"

**Fig. S1. Characterization of inflammatory colon in PI3Kδ$^{D910A}$ mice.**

(a) Hematoxylin-eosin-stained sections of WT and PI3Kδ$^{D910A}$ small intestine and colon. (b) Single enlarged C1 lymph node in PI3Kδ$^{D910A}$ mice. (c) CD25$^{+}$ Foxp3$^{-}$ precursor 1, CD25$^{-}$ Foxp3$^{+}$ precursor 2 and mature Tregs in the thymus, as a proportion of total CD4 single-positive (CD4SP) cells. n = 5, combined from 2 independent experiments. (d) Proportion of PD-1$^{+}$ cells among Tregs, CD4$^{+}$ Foxp3$^{-}$ conventional T (Tconv) cells and CD8$^{+}$ T cells in spleen, inguinal LN and separate mLN. n = 3, representative of 2 independent experiments. *, $p < 0.05$; **, $p < 0.01$, ***, $p < 0.001$; ****, $p < 0.0001$; ns, not significant.

**Fig. S2. Quantification and characterization of Tregs and conventional T cells in lymphoid tissues.**

(a) Taxonomic comparison of 16S rRNA metagenomic sequences from WT and PI3Kδ$^{D910A}$ mice. n = 6, 10-11 weeks, age-matched littermates between groups, co-housed in 3 separate cages. (b) PCA analysis of 16s rRNA sequences from fecal pellets of WT and PI3Kδ$^{D910A}$ mice (n = 6, 10-11 weeks, age-matched littermates between groups, co-housed in 3 separate cages). (c-e) Quantification of 16S rRNA sequencing reads corresponding to *Prevotella spp.* (c), *f. Clostridiales* (d) and *f. S24-7* (e). ****, $p < 0.0001$; ns, not significant.

**Fig. S3. Extended description of mass cytometric characterization of intestinal Tregs.**

(a) List of markers stained with metal-conjugated antibodies. *, markers which were used in subset gating but were excluded from Treg-specific analysis. (b) Quantification of CD4$^{+}$ Foxp3$^{+}$ Tregs as a proportion of total live CD45$^{+}$ leukocytes in intestinal tissue. n = 2. (c) Expression data of all 30 markers overlaid on UMAP visualization. Data pooled from 8 mice. SI, small intestine; IEL, intraepithelial lymphocytes; LPL, lamina propria lymphocytes.

**Fig. S4. Measurement of protein-level CCR8 expression on iTregs and ex vivo Tregs.**

(a) Proteomic analysis of Foxp3-hCD2$^+$ iTregs after 4 days in culture. n = 4 biological replicates. (b-c) Expression of CCR8 in ex vivo Tregs from cLP or spleen, measured by flow cytometry (b, representative histograms; c, quantification). n = 3 for colon, n = 6 for spleen (biological replicates).

### Fig. S1

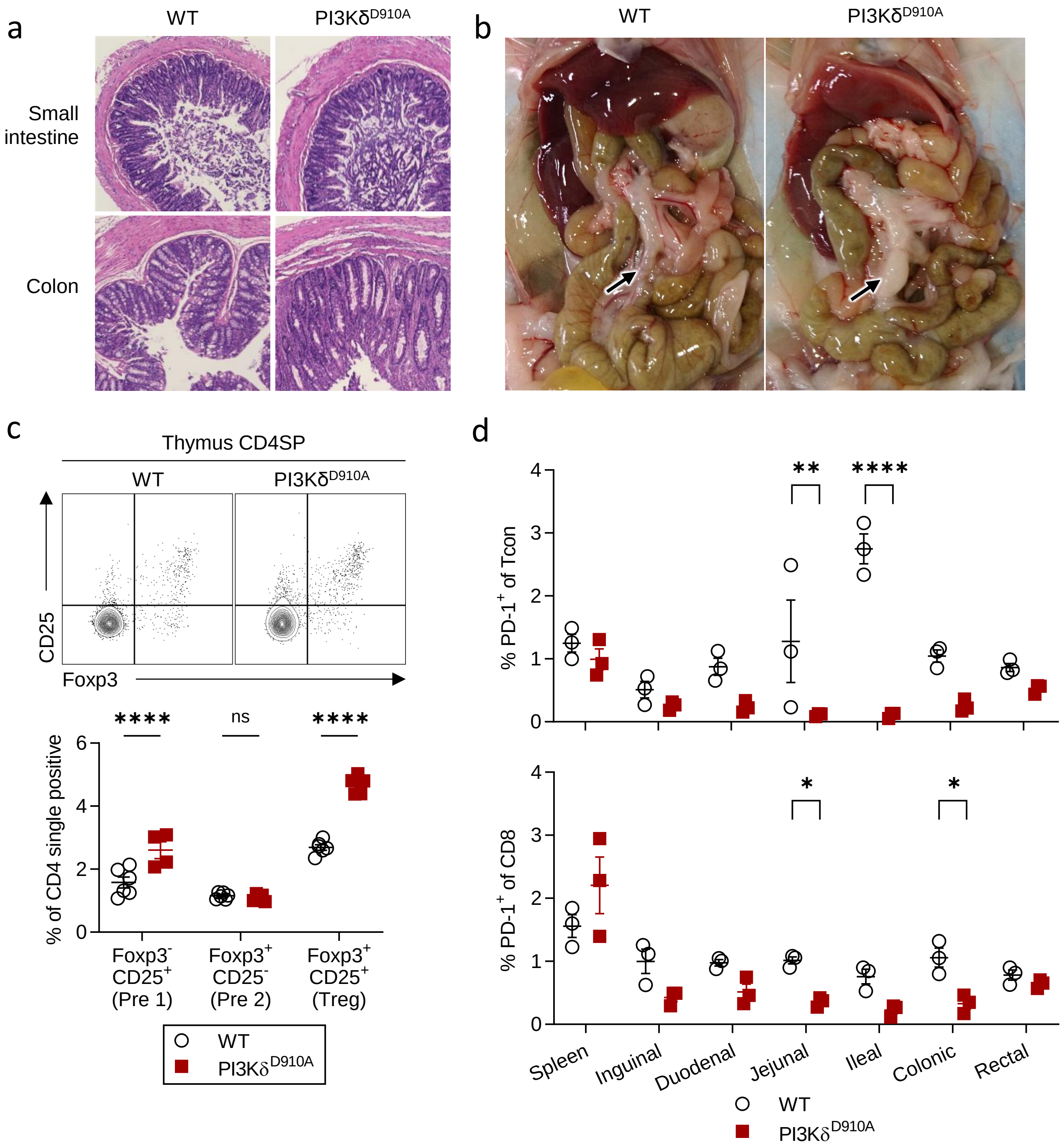

a
WT
PI3Kδ D910A
Small intestine
Colon
b
WT
PI3Kδ D910A
c
Thymus CD4SP
WT
PI3Kδ D910A
CD25
Foxp3
****
ns
****
% of CD4 single positive
Foxp3- CD25+ (Pre 1)
Foxp3+ CD25- (Pre 2)
Foxp3+ CD25+ (Treg)
WT
PI3Kδ D910A
d
% PD-1+ of Tcon
**
****
% PD-1+ of CD8
*
*
Spleen
Inguinal
Duodenal
Jejunal
Ileal
Colonic
Rectal
WT
PI3Kδ D910A

Fig. S2

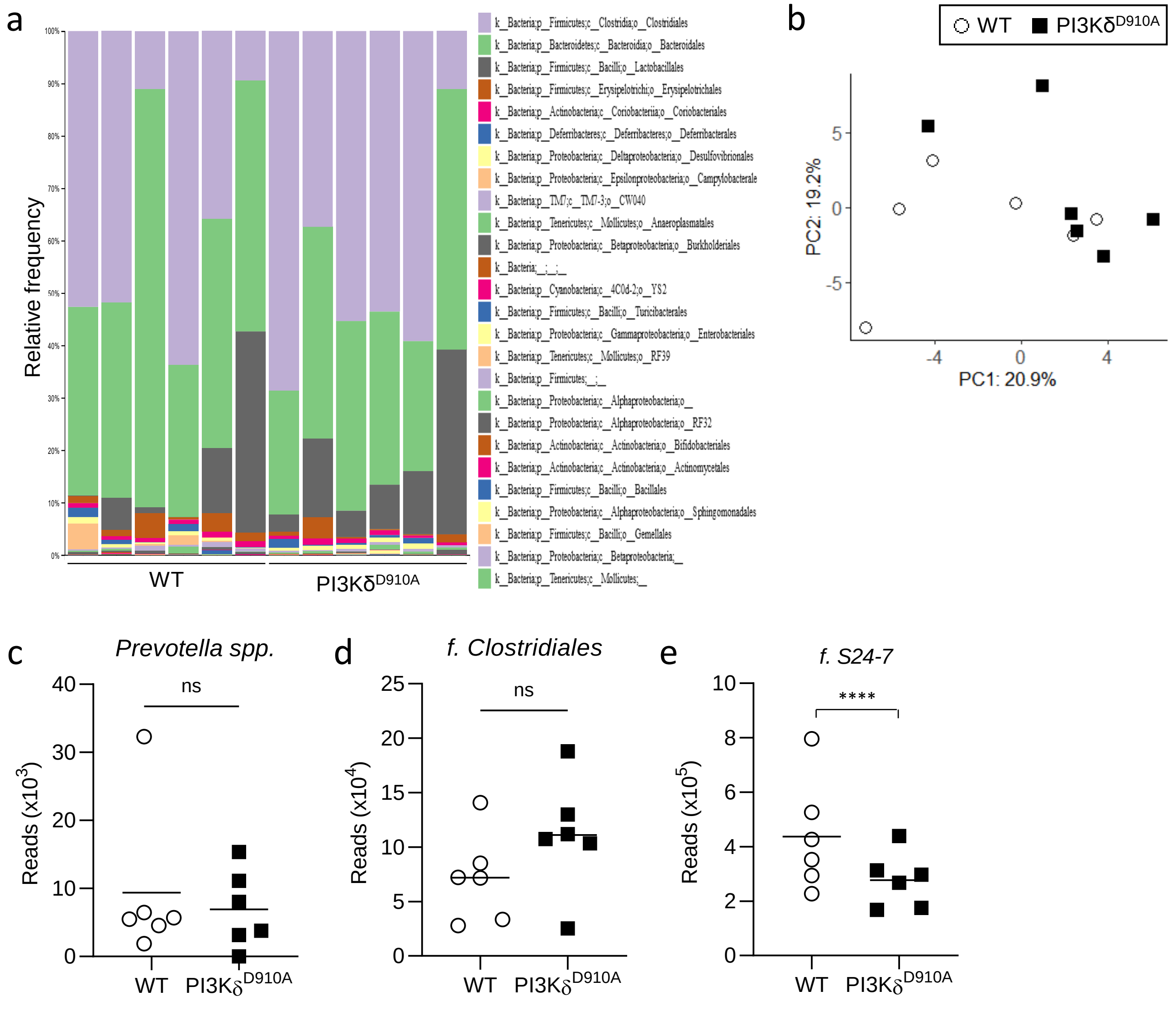

a
Relative frequency
100%
90%
80%
70%
60%
50%
40%
30%
20%
10%
0%
WT
PI3Kδ$^{D910A}$
k__Bacteria;p__Firmicutes;c__Clostridia;o__Clostridiales
k__Bacteria;p__Bacteroidetes;c__Bacteroidia;o__Bacteroidales
k__Bacteria;p__Firmicutes;c__Bacilli;o__Lactobacillales
k__Bacteria;p__Firmicutes;c__Erysipelotrichi;o__Erysipelotrichales
k__Bacteria;p__Actinobacteria;c__Coriobacteriia;o__Coriobacteriales
k__Bacteria;p__Deferribacteres;c__Deferribacteres;o__Deferribacterales
k__Bacteria;p__Proteobacteria;c__Deltaproteobacteria;o__Desulfovibrionales
k__Bacteria;p__Proteobacteria;c__Epsilonproteobacteria;o__Campylobacterale
k__Bacteria;p__TM7;c__TM7-3;o__CW040
k__Bacteria;p__Tenericutes;c__Mollicutes;o__Anaeroplasmatales
k__Bacteria;p__Proteobacteria;c__Betaproteobacteria;o__Burkholderiales
k__Bacteria;__;__;__
k__Bacteria;p__Cyanobacteria;c__4C0d-2;o__YS2
k__Bacteria;p__Firmicutes;c__Bacilli;o__Turicibacterales
k__Bacteria;p__Proteobacteria;c__Gammaproteobacteria;o__Enterobacteriales
k__Bacteria;p__Tenericutes;c__Mollicutes;o__RF39
k__Bacteria;p__Firmicutes;__;__
k__Bacteria;p__Proteobacteria;c__Alphaproteobacteria;o__
k__Bacteria;p__Proteobacteria;c__Alphaproteobacteria;o__RF32
k__Bacteria;p__Actinobacteria;c__Actinobacteria;o__Bifidobacteriales
k__Bacteria;p__Actinobacteria;c__Actinobacteria;o__Actinomycetales
k__Bacteria;p__Firmicutes;c__Bacilli;o__Bacillales
k__Bacteria;p__Proteobacteria;c__Alphaproteobacteria;o__Sphingomonadales
k__Bacteria;p__Firmicutes;c__Bacilli;o__Gemellales
k__Bacteria;p__Proteobacteria;c__Betaproteobacteria;__
k__Bacteria;p__Tenericutes;c__Mollicutes;__
b
WT
PI3Kδ$^{D910A}$
PC2: 19.2%
5
0
-5
-4
0
4
PC1: 20.9%
c
*Prevotella spp.*
ns
Reads (x10$^3$)
40
30
20
10
0
WT
PI3Kδ$^{D910A}$
d
*f. Clostridiales*
ns
Reads (x10$^4$)
25
20
15
10
5
0
WT
PI3Kδ$^{D910A}$
e
*f. S24-7*
****
Reads (x10$^5$)
10
8
6
4
2
0
WT
PI3Kδ$^{D910A}$

Fig. S3

**a**

| Lineage markers | Activation and functional markers |
|---|---|
| B220 | 4-1BB |
| BCL6 | CD103 |
| CD3 | CD25 |
| CD4 | CD38 |
| CD8 | CD39 |
| Foxp3 | CD44 |
| Gata-3 | CD5 |
| GITR | CD62L |
| Helios | CD69 |
| Ly6C | CD73 |
| RORgt | CTLA-4 |
| T-bet | ICOS |
| CD11b* | Ki67 |
| NK1.1* | KLRG1 |
| Ly6G* | Lag3 |
| TCR-β* | OX40 |
| TCR-γδ* | PD-1 |
| MHC class II* | TIM3 |

**b**

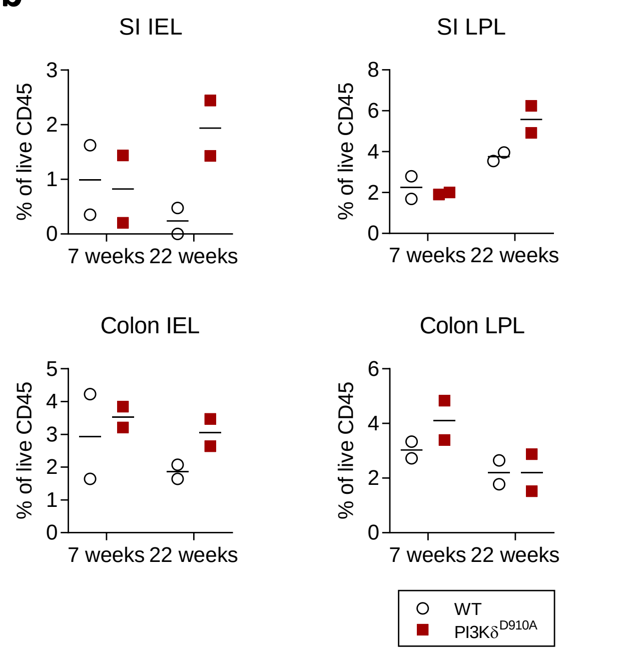


**c**

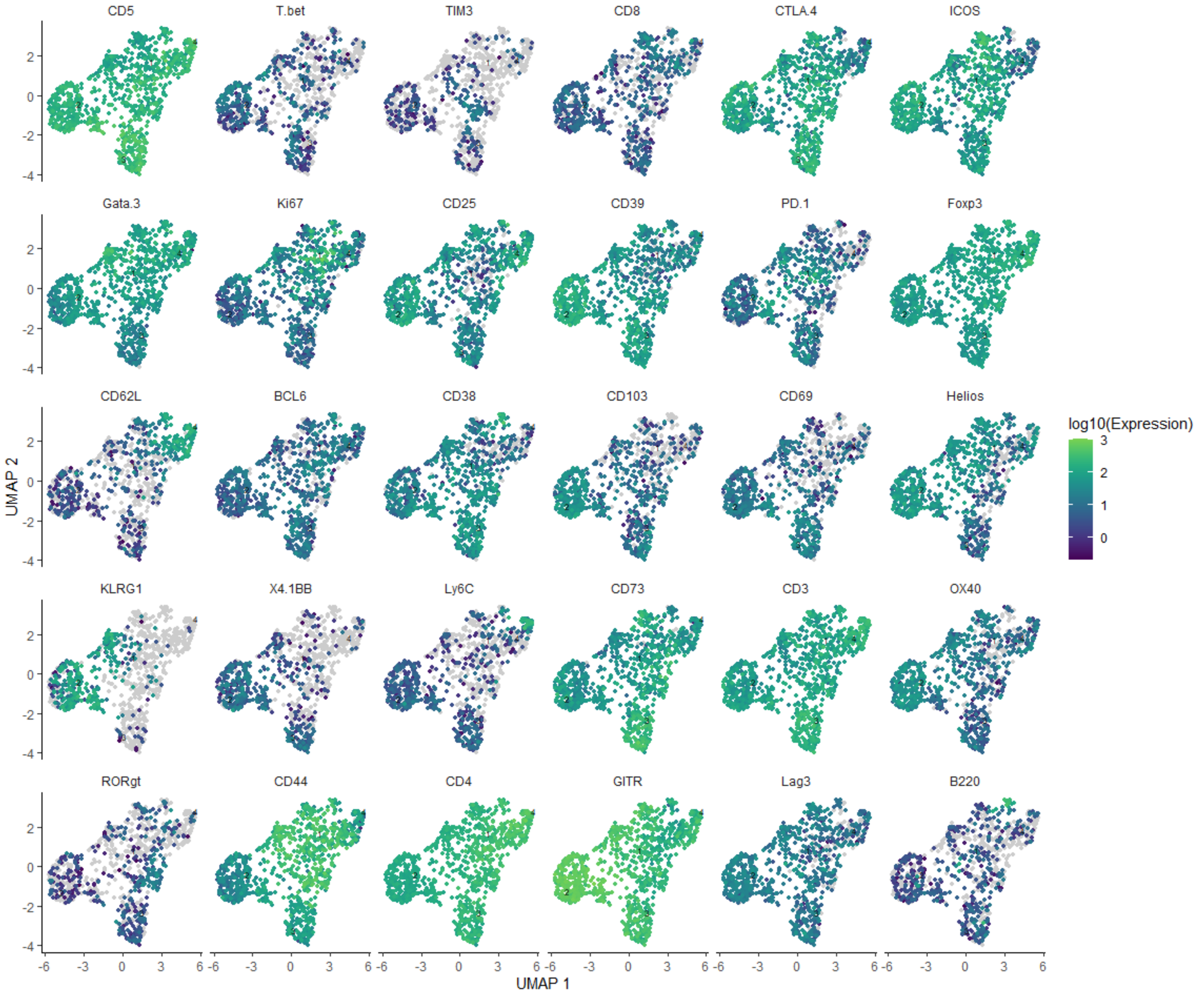

Fig. S4

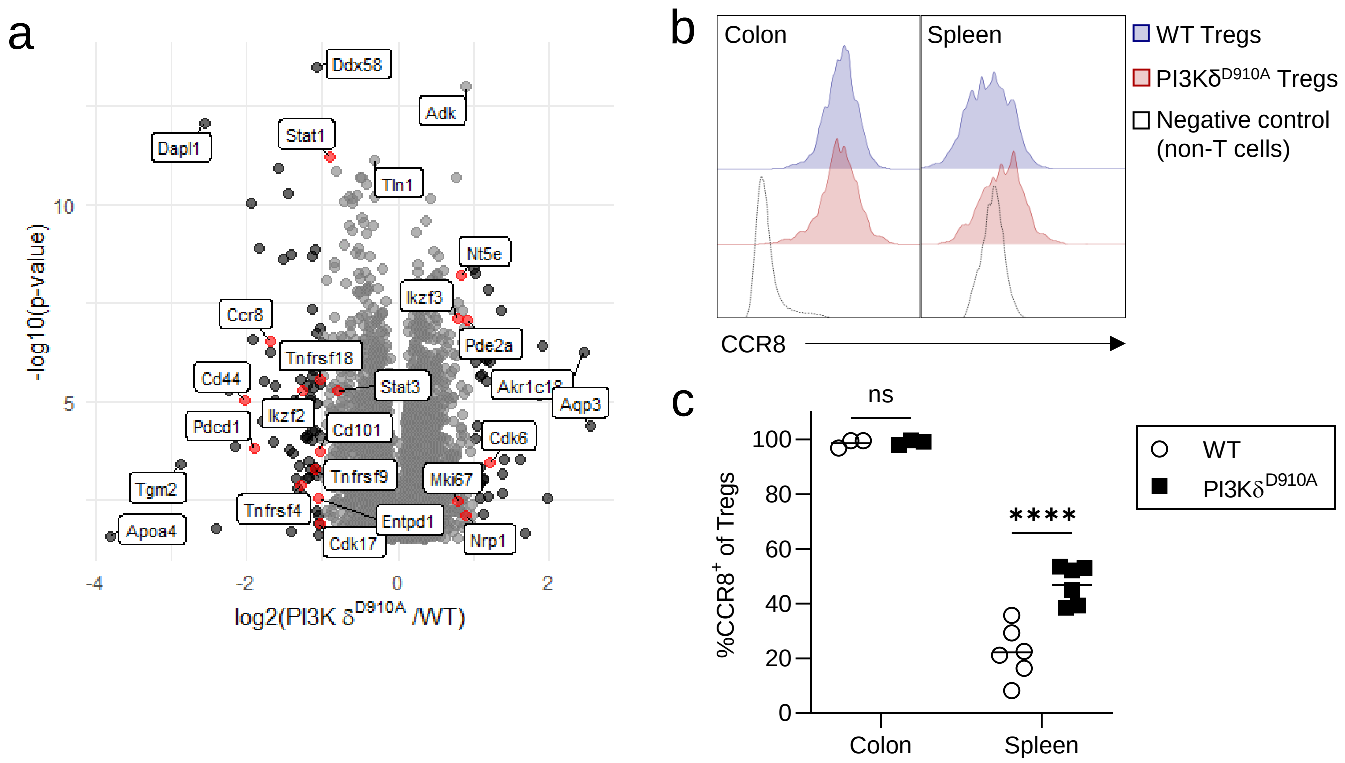

a
Ddx58
Adk
Dapl1
Stat1
Tln1
Nt5e
Ikzf3
Ccr8
Pde2a
Tnfrsf18
Cd44
Stat3
Akr1c18
Aqp3
Pdcd1
Ikzf2
Cd101
Cdk6
Tgm2
Tnfrsf9
Mki67
Tnfrsf4
Entpd1
Apoa4
Cdk17
Nrp1
-log10(p-value)
10
5
-4
-2
0
2
log2(PI3K δD910A /WT)
b
Colon
Spleen
WT Tregs
PI3KδD910A Tregs
Negative control
(non-T cells)
CCR8
c
ns
****
100
80
60
40
20
0
%CCR8+ of Tregs
Colon
Spleen
WT
PI3KδD910A
